# Large ecological benefits of small urban greening actions

**DOI:** 10.1101/2021.07.23.453468

**Authors:** Luis Mata, Amy K. Hahs, Estibaliz Palma, Anna Backstrom, Tyler King, Ashley R. Olson, Christina Renowden, Tessa R. Smith, Blythe Vogel

**Affiliations:** School of Ecosystem and Forest Sciences, University of Melbourne, Parkville 3010, Australia; School of Global, Urban and Social Studies, RMIT University, Melbourne 3000, Australia; Alfred Deakin Institute, Deakin University, Burwood 3125, Australia; School of Science, Psychology and Sport, Federation University, Churchill 3842, Australia; Office for Environmental Programs, University of Melbourne, Parkville 3010, Australia; Arthur Rylah Institute for Environmental Research, Department of Environment, Land, Water and Planning, Heidelberg 3084, Australia; School of Natural Sciences, University of Tasmania, Hobart 7005, Australia; Environmental Sustainability and Urban Design, Department of Transport, Kew 3101, Australia

**Keywords:** ecological networks, greenspaces, hierarchical models, indigenous plants, insect communities, nature in cities, urban biodiversity, urban environments, urban ecology

## Abstract

The detrimental effects of human-induced environmental change on people and other species are acutely manifested in urban environments. While urban greenspaces are known to mitigate these effects and support functionally diverse ecological communities, evidence of the ecological outcomes of urban greening remains scarce. We use a longitudinal observational design to provide empirical evidence of the putative ecological benefits of greening actions. We show how a small greening action quickly led to large positive changes in the richness, demographic dynamics, and network structure of a depauperate insect community. An increase in the diversity and complexity of the plant community led to, after only three years, a large increase in insect species richness, a greater probability of occurrence of insects within the greenspace, and a higher number and diversity of interactions between insects and plant species. We demonstrate how large ecological benefits may be derived from investing in small greening actions and how these contribute to bring indigenous species back to greenspaces where they have become rare or locally extinct. Our findings provide crucial evidence that support best practice in greenspace design and contribute to re-invigorate policies aimed at mitigating the negative impacts of urbanisation on people and other species.

## Introduction

Humans continue to cause profound, unprecedented, and accelerating changes to the function and stability of ecosystems at global scales, impacting people and other species in negative, often irreversible ways (Isbell et al. 2017; Diaz et al. 2019; IPBES 2019). Urbanisation is an acute driver of these changes, with deep eco-evolutionary effects on species occurring in or around cities (Alberti et al. 2017; Johnson and Munshi-South 2017; Palma et al. 2017; Piano et al. 2017; Merckx et al. 2018; Fenoglio et al. 2020; Mc Donald et al. 2020; Lambert et al. 2021). At local scales, however, urban greenspaces – whether large or small, permanent or temporary – are known to support functionally diverse ecological communities (Threlfall et al. 2017; Baldock et al. 2019; Mata et al. 2019; Spotwood et al. 2021), which in turn provide an array of socio-ecological benefits to urban residents (Lai et al. 2019; Mata et al. 2020; Stevenson et al. 2020). Understanding, quantifying, and managing these benefits has become a sharp focus of practitioners, professionals, and policymakers (Nilon et al. 2017; United Nations 2017; Mata et al. 2020).

Many studies highlight the positive effects of increasing vegetation structure and indigenous plant diversity on a diverse range of animal taxa in urban greenspaces (Threlfall et al. 2017; Baldock et al. 2019; Mata et al. 2021). However, there is little empirical evidence of how specific greening actions may mitigate the detrimental effects of urbanisation by facilitating the return of indigenous species that have become rare or locally extinct. Two approaches for obtaining evidence of the benefits of greening actions – and understanding what the ecological outcomes would have been if actions had not taken place – are experimental ‘randomised controlled trials’ and counterfactual ‘before-after-control-impact’ evaluations (Christie et al. 2019). Presently, however, applying these across large scales remain largely unfeasible, due to cost, logistics, and project duration constraints. Indeed, the few studies reporting ecological benefits of greening have used instead ‘space-for-time substitutions’ (De Palma et al. 2018), comparing outcomes of sites that have been greened for some years with non-greened controls (Archibald et al. 2017; Mody et al. 2020).

As we understand, no study to date has sought to track how the putative ecological benefits accrued across the lifespan of specific greening actions using a longitudinal observational design. This approach has the advantage that the ecological state of the system is known before the actions occur, as opposed to space-for-time substitutions, where the baseline state is assumed or inferred (De Palma et al. 2018). Most relevant to urban areas, the approach is ideally suited for opportunistic studies, where researchers are made aware of the execution of the greening actions with short notice and there is no availability of matching control sites.

Here, we report empirical evidence of the ecological benefits of urban greening. We collected a plant-insect interactions dataset before, and for three years after, a greenspace received a small greening action within a densely urbanised municipality. We then assessed how (1) insect species richness; (2) the probabilities of occurrence, survival and colonisation of the insect community; and (3) the plant-insect network structure varied across the four years of the study. To complement traditional analyses focusing on species richness, our analytical approach was designed to forecast probability statements about demographic rates (Kéry and Schaub 2012) and harness theoretical advances in network science (Kaiser-Bunbury and Blüthgen 2015; Tylianakis and Morris 2017; Guimarães 2020). As such, we provide a foundation to demonstrate whether ecological communities in greened urban sites are developing on trajectories towards robust and resilient states. Most importantly, our study contributes critical evidence-base to support future greening projects and the practice, policy, and decision-making for protecting nature in urban environments (Mata et al. 2020).

## Materials and methods

### Study site

Our study was conducted across four years (2016-19) at the Tunnerminnerwait and Maulboyheenner memorial site, a small (195 m2) greenspace in the City of Melbourne, Victoria, Australia (Fig. S1). The site is adjacent to a major road, surrounded by large buildings, and embedded in a dense urban matrix (Fig. S2). The site’s vegetation prior to March 2016 was limited to a kikuyu (*Cenchrus clandestinus*) lawn and two spotted gum (Corymbia maculata) trees (Table S1). In mid-April 2016, 80% of the site was substantially transformed through weeding, addition of new topsoil, soil decompaction and fertilisation, organic mulching, and the addition of 12 indigenous plant species (Table S1; Fig. S3). In between 2017 and 2018, one plant species was added and four perished (Table S2). After that, the composition of the plant community remained stable.

### Data collection

We conducted 14 insect surveys across four years – four before the greening actions in 2016 (henceforth Year 0), four in 2017 (Year 1), three in 2018 (Year 2), and three in 2019 (Year 3). Surveys were conducted between late-January and early-April each year. We used an entomological net to sample each plant species occurring at the site for ants, bees, beetles, flies, hemipteran bugs, and wasps. The number of sweeps per plant was standardised as a proportion of the species’ volume within the site (Mata et al. 2021). Specimens were identified to species or morphospecies (henceforth species), assigned to functional groups (detritivores, herbivores, predators, or parasitoids), and grouped by main evolutionary clade (Table S3).

### Modelling species richness

We used a variation of the hierarchical metacommunity model (Kéry and Royle 2016) described by Mata and colleagues (2021) to assess how the species richness of indigenous insect species varied across years. ‘Plant species’ was the unit of analysis for drawing inferences on insect species occupancy and the repeated temporal samplings constituted the unit of detection replication. The model is structured around three levels: the first one models species occupancy; the second species detectability; and the third treats the occupancy and detection parameters for each species as random effects (Kéry and Royle 2016).

We specified the occupancy level model as:

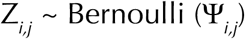

where Ψ_*i,j*_ is the probability that insect species *i* occurs at plant species *j*, and the detection level model as:

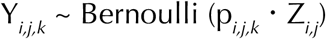

where p_*i,j,k*_ is the detection probability of insect species *i* at plant species *j* at temporal replicate *k*.

The occupancy and detection level linear predictors were specified on the logit-probability scale as:

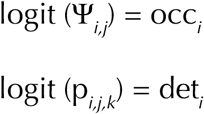

where occ_*i*_ and det_*i*_ are the species-specific random effects, which were specified as:

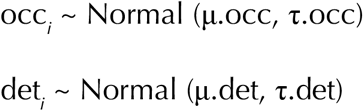

where the metacommunity mean occupancy (μ.occ) and detection (μ.det) hyperpriors were specified as Uniform (0, 1) and the metacommunity precision occupancy (τ.occ) and detection (τ.det) hyperpriors as Gamma (0.1, 0.1).

We then used the latent occurrence matrix Z_*ij*_ to estimate the insect species richness associated with each plant species SR*j* through the summation:

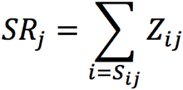

where S_*ij*_ is a ‘specificity’ vector indexing the insect species to be included in each plant species′ estimate (Mata et al. 2021). SR_*j*_ is then an estimate that accounts for plant-insect specificity, in which, for each plant species, the observed insect species are included with probability of occurrence = 1 and a limited random sub-sample of other insect species occurring in the study area are included with their 0 < Z < 1 estimated probabilities of occurrence. This makes it possible to work within the reasonable ecological assumption that at the study site not every insect species will be associated with every co-occurring plant species.

As these calculations were conducted within a Bayesian inference framework, the insect species per plant species estimates were derived with their full associated uncertainties. We ran individual models for each year and used the models’ occurrence matrices to estimate the insect species richness associated with each plant species. For each year, we averaged the species richness estimates of the (1) ‘baseline’ plant species that were present in Year 0 and (2) ‘greening action’ plant species that were added in or after Year 1 to obtain posterior distributions for each plant group that could be statistically compared within and across years.

### Modelling occupancy and demographic rates

We used a multiseason site-occupancy model (Kéry and Schaub 2012) to assess how the probabilities of occurrence, survival, and colonisation of the insect community varied across years. The model is equivalent to a metapopulation model, where changes between year *t* and year *t*+1 in insect occupancy are expressed as a function of the probabilities of colonisation on plant species unoccupied in year *t*, and of survival on plant species occupied in year *t*. As for the metacommunity model, ‘plant species’ was the unit of analysis and the repeated temporal samplings constituted the unit of detection replication. The model is structured around two levels: one for occupancy and a second one for detectability (Kéry and Schaub 2012).

We specified the occupancy level model in the baseline year (Year 0) as:

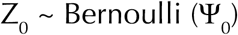

where Ψ_0_ is the probability of occurrence in Year 0.

We specified the occupancy level models for subsequent years (Year 1, Year 2, and Year 3) as

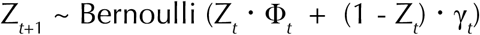

where Φ and γ are the probability of colonisation and probability of survival, respectively.

We then calculate the probability of occurrence for Year 1, Year 2, and Year 2 as derived quantities with the following equations:

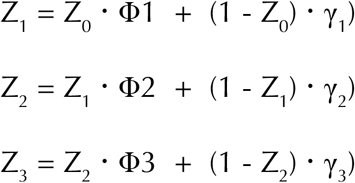

Lastly, we specified the detection level model as:

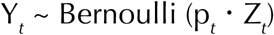

where p_*t*_ is the probability of detection.

All occurrence, colonisation, survival, and detection priors were specified as Uniform (0, 1).

### Modelling network metrics

To assess how network structure varied across years, we first organised the data into plant species by insect clade matrices (one for each of the 14 replicated surveys), with cell values representing the number of times insect species within a given clade were recorded interacting with each plant species. We then used the matrices to calculate four network-level (interaction richness, interaction diversity, interaction evenness, and network specialisation *H*_2_′) and two species-level (plant *d*′_plants_ and insect *d*′_insects_ specialisation) metrics. All metrics were calculated with the R package bipartite (Dormann et al. 2008). Lastly, we used generalised linear models to estimate how network metrics varied across years. All models were structured around a single level, in which the model for the given network metric was specified as

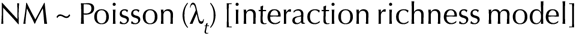

or

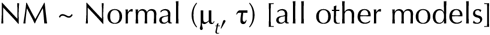

where the expected counts λ_*t*_ and means μ_*t*_ for each year were given Normal (0, 0.001) priors and

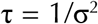

where σ was given a Uniform (0, 100) prior.

### Bayesian inference implementation

We estimated all model parameters under Bayesian inference, using Markov Chain Monte Carlo (MCMC) simulations to draw samples from the parameters′ posterior distributions. We implemented models in JAGS (Plummer 2003), as accessed through the R package jagsUI (Kellner 2016). We used three chains of 2,500 iterations, discarding the first 500 in each chain as burn-in. We visually inspected the MCMC chains and the values of the Gelman-Rubin statistic to verify acceptable convergence levels of R-hat < 1.1 (Gelman and Hill 2007).

## Results

Overall, we recorded 94 insect species, representing 22 detritivore, 35 herbivore, 11 predator, and 26 parasitoid species (Tables S3-S4). As many as 97% of all recorded species were indigenous to the study area. The most commonly occurring species was the minute brown scavenger beetle *Cortinicara sp. 1* (Cucujoidea: Latridiidae), accounting for 15% of all records.

### Species richness

We found that after only one year the twelve plant species that were planted during the greening actions supported an estimated 4.9 times more insect species than the two plant species comprising the pre-greening vegetation on site (Table S5; Fig. 1a). By year three, the nine remaining plant species supported an estimated 7.3 times more insect species than the baseline plant species (Table S5; Fig. 1a). We also found marked within-year statistical differences in the number of insect species per plant species, with the average greening action plant species showing 1.6, 2.7, and 4.2 times more insect species in year 1, year 2, and year 3, respectively, than the average baseline plant species for the same year (Table S5; Fig. 1a).

**Fig 1.**
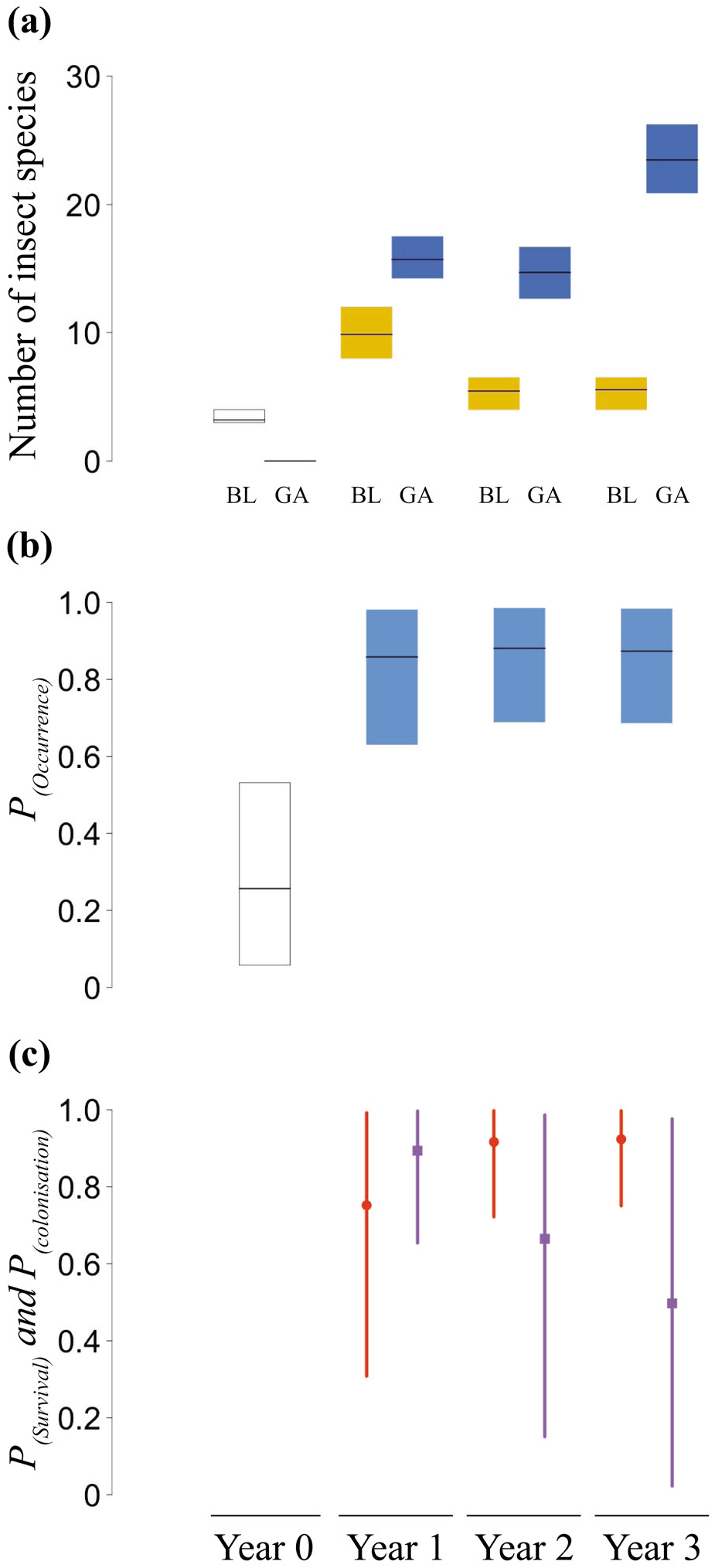
(a) Number of insect species by year, as estimated by the hierarchical metacommunity model. Yellow and blue boxes represent baseline (BL) and greening action (GA) plant species, respectively. (b) Probability of occurrence and (c) probabilities of survival (red) and colonisation (purple), as estimated by the multiseason site-occupancy model. In (a) and (b), the horizontal black lines represent the mean response, and the boxes the uncertainty associated with 95% Credible Interval. In (c), the circle (survival) and square (colonisation) represent the mean response, and the vertical lines the uncertainty associated with 95% Credible Interval.

The marked statistical differences we found after one, two, and three years in the estimated number of insect species between the greening action and baseline plant species were consistent across all functional groups (Table S5; Fig. 2). However, while the mean number of herbivores and parasitoids estimated for the greening action plant species was higher in year 3 than in year 1, the 95% CI for year 3 in these groups slightly overlaps that of year 1 (Table S5; Fig. 2). We also found that the number of estimated predators between the greening action and baseline plant species was only statistically different in year 3 (Table S5; Fig. 2).

**Figure 2.**
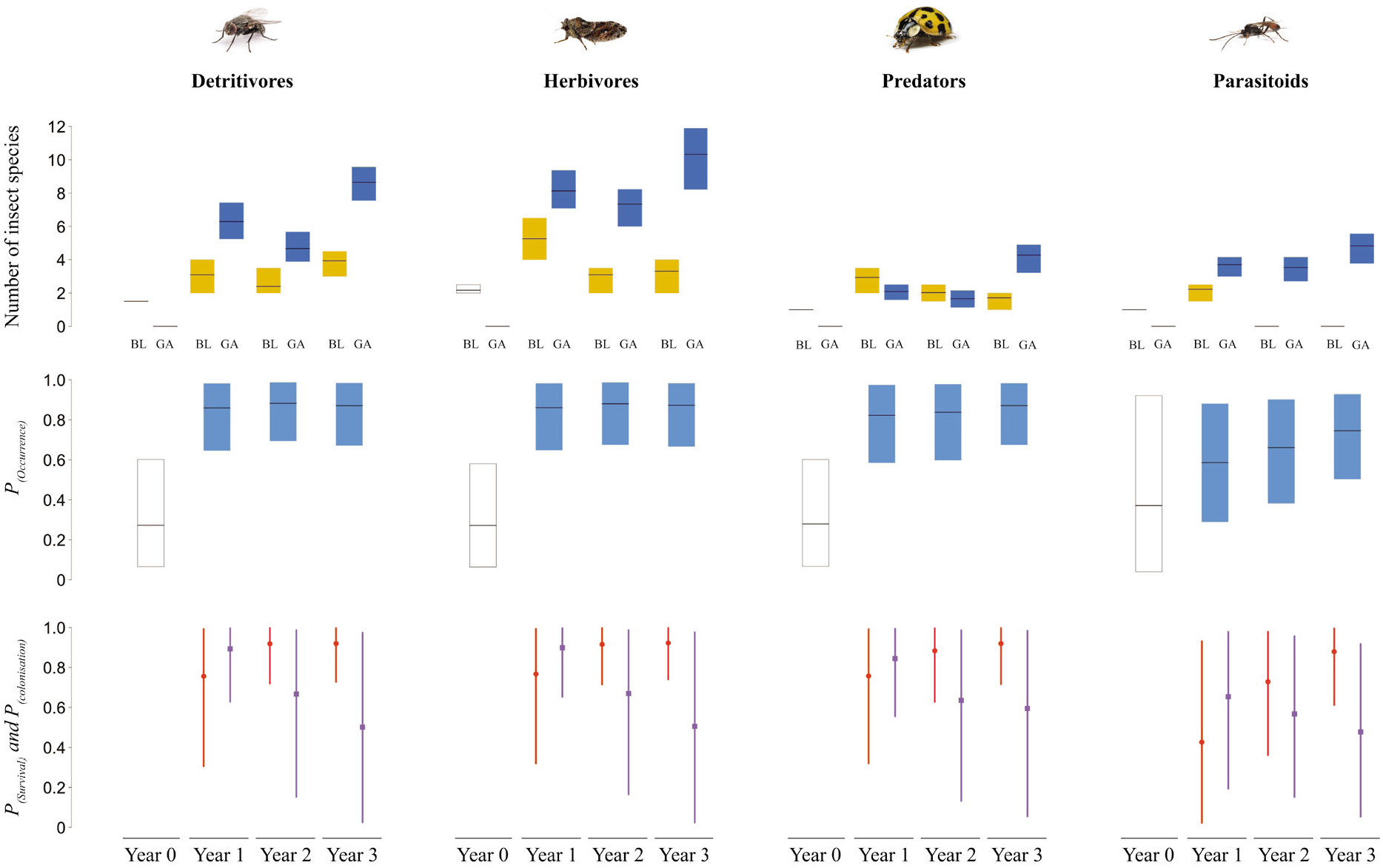
Number of insect species, and probabilities of occurrence, survival and colonisation, by year, for detritivore, herbivore, predator and parasitoid insect species. Top row: number of insect species by year, as estimated by the hierarchical metacommunity model. Yellow and blue boxes represent baseline (BL) and greening action (GA) plant species, respectively. Middle row: probability of occurrence for Year 0 (white box) and Year 1, Year 2, and Year 3 (blue boxes), as estimated by the multiseason site-occupancy model. Bottom row: probabilities of survival (red) and colonisation (purple), as estimated by the multiseason site-occupancy model. Horizontal black lines, circles and squares represent mean response, and boxes and vertical lines the uncertainty associated with 95% Credible Interval.

### Occupancy, survival, and colonisation

We found a marked statistical difference for the probability of occurrence of insects between the baseline and greening action years, with model estimates showing a 3.4-fold increase in the mean probability of occurrence of insects from year 0 to year 3 (Table S6; Fig. 1b). The demographic dynamics of the insect community in year 1 were predominantly driven by colonisation (Table S6; Fig. 1c). By year 2, dynamics diametrically shifted to a system predominantly driven by survival, a trend that was only slightly more pronounced in year 3 (Table S6; Fig. 1c).

These patterns for the whole insect community were consistent across all functional groups (Table S6; Fig. 2). However, we are unable to statistically compare the probability of occurrence of parasitoids between the baseline and greening actions years (Table S6; Fig. 2), as the model estimate for year 0 is based on a single observation, and therefore we only have a point estimate for this year, with no uncertainty associated to it.

### Network metrics

We found that the structure of the plant-insect network varied substantially across the four years of the study (Fig. 3a). Three years after greening, the number and diversity of interactions was, on average, 14.6 and 3.4 times higher, respectively, in the greened compared to the baseline network (Table S7; Fig. 3b). We uncovered, however, no statistical differences in interaction evenness between baseline and greened networks (Table S7; Fig. 3b).

**Figure 3.**
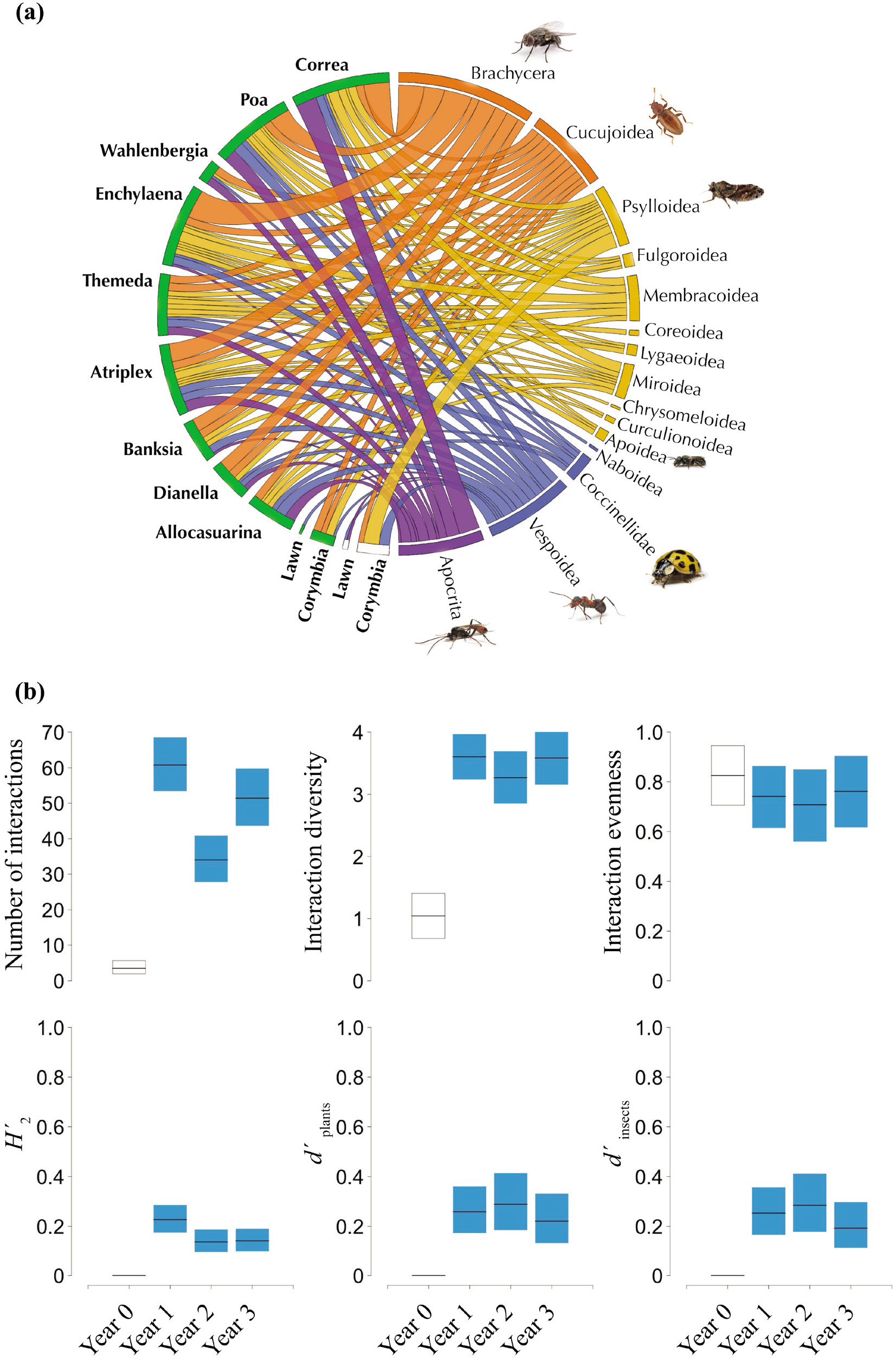
(a) Bipartite quantitative networks of interactions (chords) between plant species (baseline: white boxes; greening (year 3): green) and insect clades (detritivores: orange; herbivores: yellow; predators: blue; parasitoids: purple). Chord width reflects the relative richness of insect species from the given clade represented in the interaction. (b) Network metrics in Year 0 (white box) and Year 1, Year 2, and Year 3 (blue boxes), as estimated by the generalised linear models. Metrics include number of interactions, interaction diversity, interaction evenness, network specialisation (*H*_*2*_′), and plant (*d*′_plants_) and insect (*d*′_insects_) specialisation. Horizontal black lines represent mean response and boxes the uncertainty associated with 95% Credible Interval.

We found that specialisation (*H*_*2*_′) was consistently low across years in the greened network, a pattern that was paralleled by the metrics of plant (*d*′_plants_) and insect (*d*′_insects_) specialisation (Table S7; Fig. 3b). The low number of interacting species in year 0 prevented us from calculating these metrics for the baseline network.

## Discussion

In this study, we provide robust empirical evidence of the ecological benefits of urban greening. We show how a small greening action conducted in a densely urbanised area led to large positive changes in the richness, demographic dynamics, and network structure of a depauperate insect community. An increase in the diversity and complexity of the plant community led to, after only three years, a large increase in insect species richness, a greater probability of occurrence of insects within the greenspace, and a higher number and diversity of interactions between insects and plant species. Our findings therefore demonstrate that large ecological benefits may be derived from investing in small urban greening actions and that these may bring indigenous insect species back to urban areas where they have become rare or locally extinct. This is crucial evidence to help support local, regional, and global policy aimed at mitigating the negative impacts of urbanisation for the benefit of people and other species.

Our findings are consistent with studies documenting the positive effects of vegetation complexity and indigenous plant species on the biodiversity of insects and other animal taxa in urban environments (Threlfall et al. 2017; Mata et al. 2021). Indeed, greening actions present an excellent opportunity to bring indigenous plant species back into our cities and towns (Mata et al. 2020). Increasing the distribution and cover of indigenous plants across cities may in turn contribute to mitigate the undesired consequences of nonnative plant introductions (Pyšek et al. 2020). Furthermore, indigenous plants may provide optimal options to greenspace managers seeking to select plant species that are well-adapted to local climates and that contribute to more natural, weed reducing leaf litters (Mody et al. 2020).

Our results provide compelling evidence that colonisation was the primary demographic process driving the large increases in insect richness observed one year after the greening action. From year two onwards, colonisation is replaced by survival as the system’s main demographic process. These observations demonstrate that greening actions can bring about positive ecological changes within a few years after implementation. Our findings have, therefore, important implications for policymakers seeking to incentivise stakeholders to uptake, fund, and maintain greening actions across cities.

The sharp rise in insect richness and community-level probability of occurrence was mirrored by an equally sharp rise in the number and diversity of plant-insect interactions. This concurs with Kaiser-Bunbury and colleagues (2017) who demonstrated the positive effects of ecological restoration on disturbed plant-pollinator communities. Another exciting parallel between our studies is the low levels of network (*H*_*2*_′) and species (*d*′) specialisation reported in restored/greened networks, indicating higher functional redundancy and lower mutual dependencies (Kaiser-Bunbury and Blüthgen 2015). These findings demonstrate the key role that restoration and greening actions can play in boosting the resilience of disturbed ecosystems, while facilitating their functional robustness to local species loss.

Our approach to quantifying how greening actions drive positive ecological changes has allowed us to expand current applied research practice to a more intricate exploration of demographic dynamics and network structure across multiple trophic levels using a longitudinal observational design. This was possible through a reproducibly, multi-year data collection protocol and an analytical approach that is supported by recent advances in hierarchical modelling and ecological network science. We hope that our study will serve as a catalyst for a new way to demonstrate the ecological benefits of urban greening.

The flexible methodology we present here can be adapted to include multiple sites, seasons, longer time series, matching control sites, and other functional groups such as pollinators and frugivores. Comparing actions carried out in greenspaces of varying sizes could shed light into area-dependent responses – community and network dynamics that are only manifested beyond a minimum greenspace size. For example, can small greenspaces be designed or modified to provide the levels of food availability and resource heterogeneity required to promote bird colonisation and survival, and, therefore, buffer the city-wide metacommunity against extinction? The prospect of being able to answer this and other related questions serves as a stimulus for future research.

Our findings provide much needed scientific evidence that demonstrate how simple greening actions have real, quantifiable effects on the richness, demographic dynamics, and network structure of complex ecological communities. This understanding is fundamental to assist architects, engineers, developers, and planners design greenspaces that serve people and other species. This is particularly important given the immense value of greenspace in cities. Crucially, our findings set robust pathways for greening projects to support evidence-based practice and policy, therefore supporting decision-makers in charge of protecting and bringing back nature into urban environments.

## Acknowledgements

The authors acknowledge the Traditional Custodians of the land and waterways on which the project took place and where we work and live, the Wurundjeri and Bunurong people of the Kulin Nations, and the muwinina people of lutruwita (Tasmania), the Country of Tunnerminnerwait and Maulboyheenner. We pay our respects to their Elders, past, present and emerging, and honour their deep spiritual, cultural, and customary connections to the land. We would like to extend our heartfelt thanks Zena Cumpston, Maddison Miller, and Jade Kennedy for sharing with us their passion and knowledge of Australian Indigenous culture and for helping us recognise and appreciate the cultural significance of indigenous plant species. The authors would also like to sincerely thank Yvonne Lynch, Lingna Zhang, and Ian Shears at the City of Melbourne for providing access to and data on the Tunnerminnerwait and Maulboyheenner memorial site. Thanks to Udo Schimdt and Bernard Dupont for the Cucujoidea and Psylloidea images, and to Circos for the software to produce the chord diagram. LM would like to thank Marc Kéry for providing advice and inspiration on hierarchical modelling and acknowledges financial support from the Australian Government’s National Environmental Science Program through the Clean Air and Urban Landscapes Hub during the conceptualisation and data collection period of the study.

## Data availability statement

Data and codes to reproduce models and plots are already published and publicly available in Zenodo: https://zenodo.org/record/5140619

