## Supplementary Information for "Large ecological benefits of small urban greening actions"

**Large positive ecological changes of small urban greening actions**

Supplementary Information

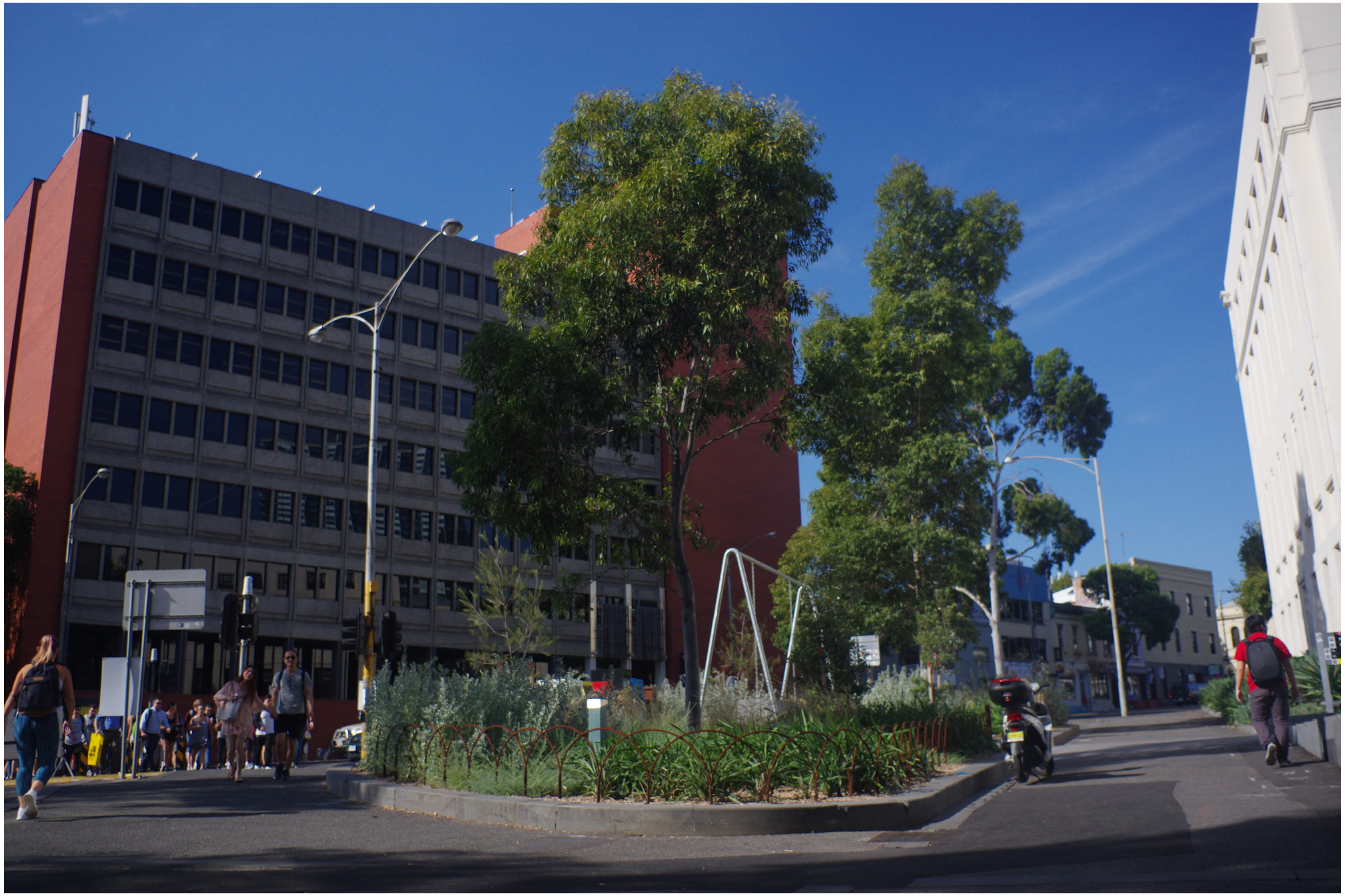

**Figure S1**. The Tunnerminnerwait and Maulboyheenner memorial is located at the intersection of Franklin Street and Victoria Street in the Melbourne Central Business District (City of Melbourne, Victoria, Australia). The site hosts the 'Standing by Tunnerminnerwait and Maulboyheenner' marker by artists Brook Andrew and Trent Walter, which commemorates the lives of Tunnerminnerwait and Maulboyheenner, two Tasmanian Aboriginal men who were publicly hanged in the vicinity of the site in 1842. Photo from March 2017 (Photo credit: Luis Mata).

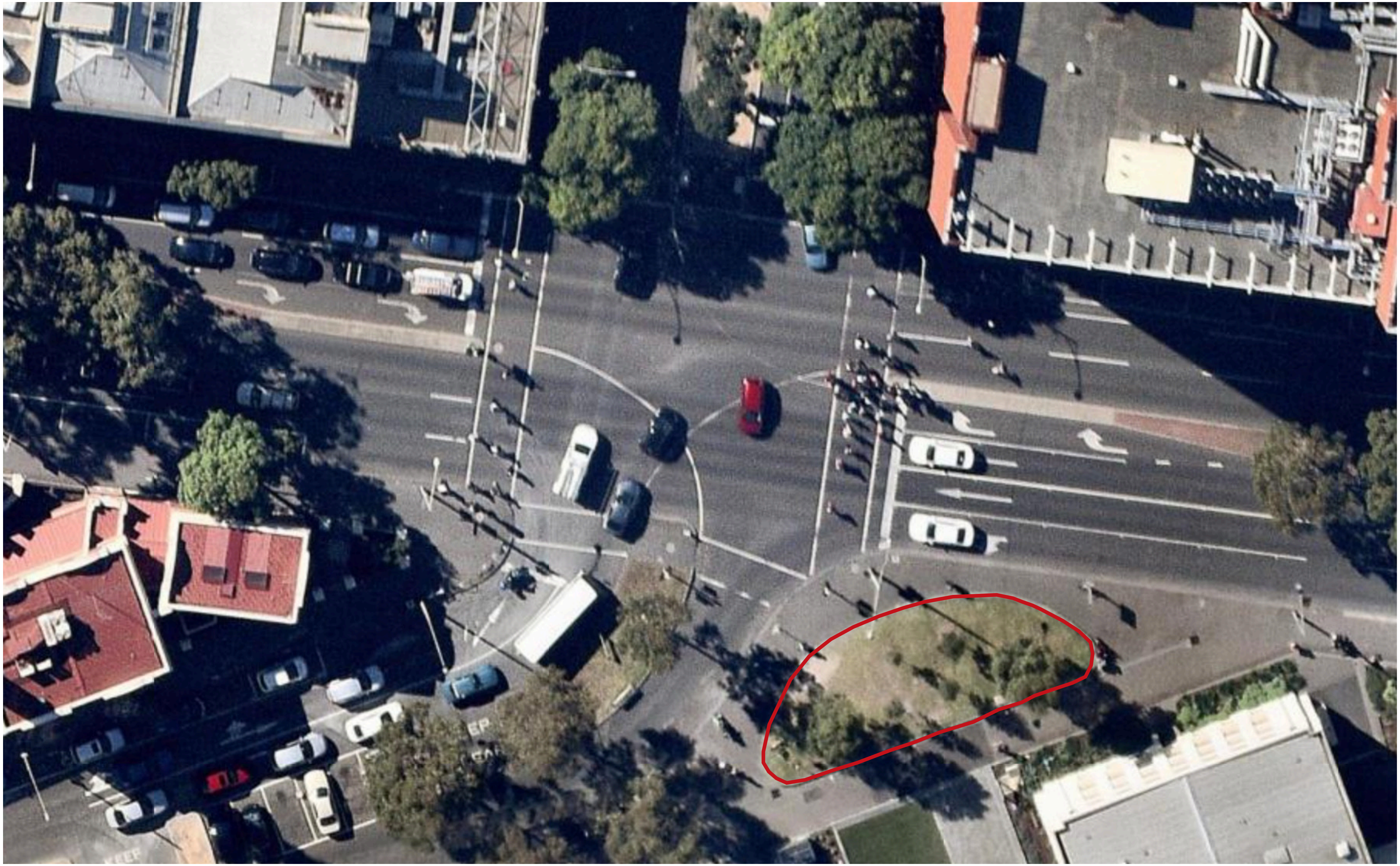

**Figure S2**. Aerial image of the Tunnerminnerwait and Maulboyheenner site (red outline) and its surroundings. Photo from March 2016 (Photo credit: Google Earth via Nearmap).

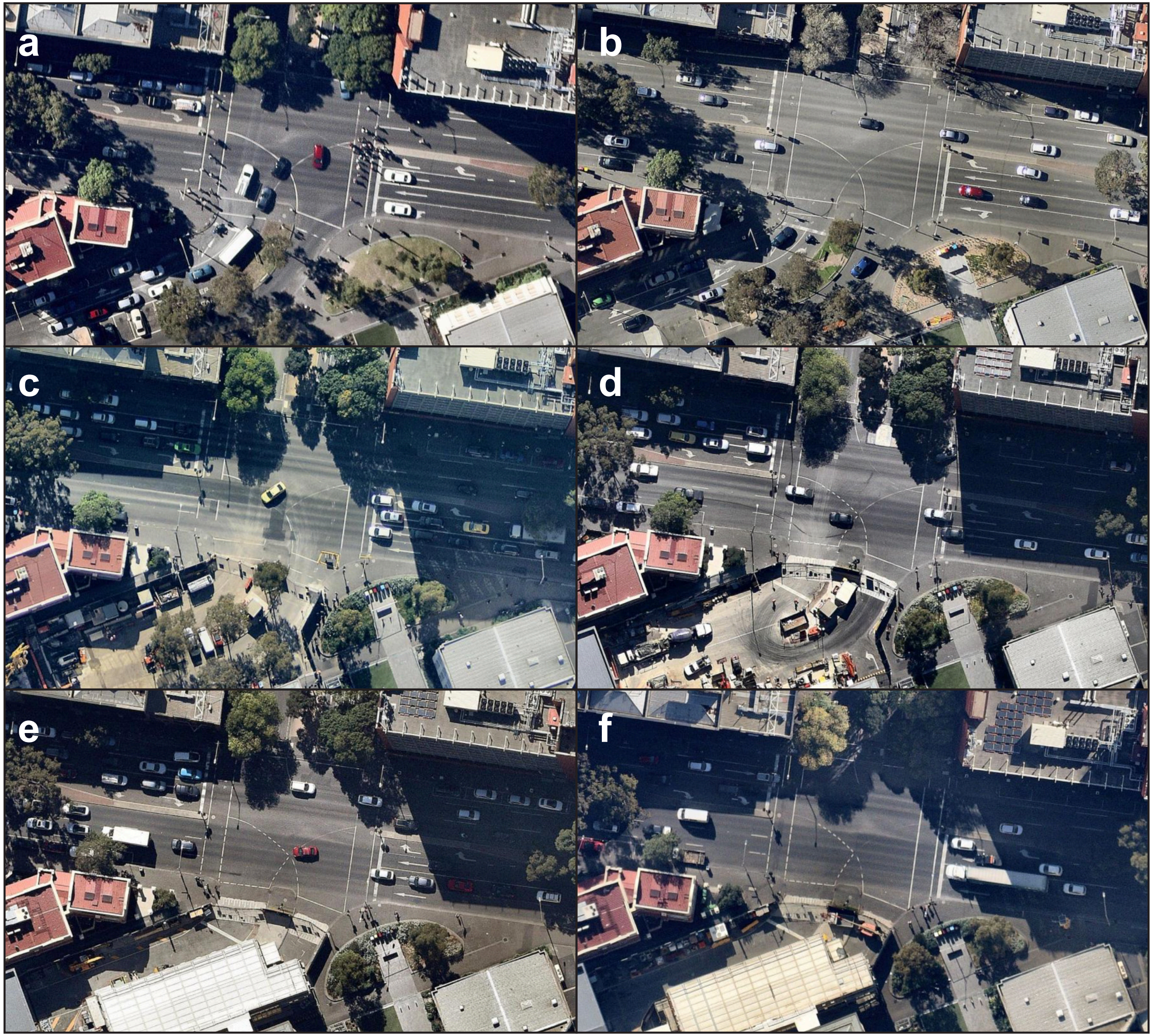


**Figure S3** Aerial images of the Tunnerminnerwait and Maulboyheenner from before (**a**) and after the site was greened (**b-f**). Photos from March 2016 (**a**), October 2016 (**b**), May 2017 (**c**), April 2018 (**d**), April 2019 (**e**), and April 2021 (**f**). The site's exact location and boundaries is shown in Figure S2. (Photo credits: Google Earth via Nearmap).

**Table S1**. Detailed description of the 15 plant species that were part of the study.

| Common name | Species | Family | Growth form | Origin | Year 0 [2016] | Year 1 [2017] | Year 2 [2018] | Year 3 [2019] | IUCN cat. | General description | Distribution |
|---|---|---|---|---|---|---|---|---|---|---|---|
| Black sheoak | *Allocasuarina littoralis* | Casuarinaceae | Tree | Locally indigenous | | | | | VU | Small upright tree to 8 m tall with foliage of mainly ascending fine branchlets and leaf teeth not overlapping. Branchlets superficially resemble pine needles. Bark is fissured and on old trees dark and deeply furrowed. Plants dioecious. Female flowers reddish, cones cylindrical usually longer than broad. Male flower spikes dark brown. Flowers Mar-Jun. | Common throughout South-east Victoria in a range of woodland habitats. Mainly growing on near-coastal sands but also on heavy clay soils or among rocks. |
| Coast saltbush | *Atriplex cinerea* | Amaranthaceae | Shrub | Locally indigenous | | | | | | Dense, spreading shrub (1-2 m tall, 2-3m wide) with brittle branches and silver-grey ovate to oblong leaves to 8 cm long. Plants dioecious. Male flowers reddish-purple, in dense, globular clusters on spikes; female flowers are cream and sit in the leaf axils. Flowers Oct-Jan. | Requires well drained sandy soil on shorelines above high tide levels in full sun. Occurs throughout Australia in coastal areas. |
| Coast banksia | *Banksia integrifolia* | Proteaceae | Tree | Locally indigenous | | | | | EN | Open, erect and spreading tree up to 20 m tall with thick tessellated bark. Leaves dark green above and silver-velvety below, up to 15 cm long and 3.5 cm wide, margins on juvenile leaves may be toothed, but adult leaves entire. Pale yellow flowers are borne on brushes to 12 cm long. Flowers Feb-Sep. | Requires well drained soils and full sun. Occurs on the coastal margin of South-east Victoria and along the East coast of Australia in a range of habitats from coastal dunes to mountains. |
| Kikuyu | *Cenchrus clandestinus* | Poaceae | Graminoid | Introduced | | | | | | A rhizomatous grass with matted roots and a grass-like or herbaceous habit. The leaves are green, flattened or upwardly folded along the midrib, 10-150 mm long, and 1-5 mm wide. The apex of the leaf blade is obtuse. Sheaths are ciliate along margins. Inflorescence remaining wholly or largely enclosed within uppermost leaf-sheath. Stamens briefly but prominently exserted at anthesis, the fine filaments 2-4 cm long. Flowers Jan-Apr. | Naturalised within Victoria, this species prefers sandy soils of lowland areas, and has failed to thrive in very heavy soils, dry areas and cooler montane zones. Widely used as a pasture and lawn species. Kikuyu invades dry coastal vegetation, heathland and heathy woodland, lowland grassland and grassy woodland, dry sclerophyll forest and woodland, and riparian vegetation. |
| Prickly currant-bush | *Coprosma quadrifida* | Rubiaceae | Shrub | Locally indigenous | | | | | NT | Open, upright, spiny shrub 2 - 4 m tall. Foliage thin, variable leaf shape - lanceolate to broad ovate, conspicuous veins below. Plants dioecious. Inconspicuous greenish flowers, terminal on branchlets. Fruit, small bright red berries. Flowers Jan-Mar. | Widespread throughout South-east Victoria. Often common in moist open forests, riparian scrub, fern gullies and rainforest. |
| White correa | *Correa alba* | Rutaceae | Shrub | Locally indigenous | | | | | EN | Dense, spreading shrub to 2 m tall. Thick, elliptic, grey-green leaves, sparsely tomentose above, paler and densely velvety below. Flowers white, open and 4-dentate (petal-like), not pendulouse bells. Flowers Sep-Feb. | Common in near-coastal heaths and woodlands. |

**Table S1 (Cont.)**. Detailed description of the 15 plant species that were part of the study.

| Common name | Species | Family | Growth form | Origin | Year 0 [2016] | Year 1 [2017] | Year 2 [2018] | Year 3 [2019] | IUCN cat. | General description | Distribution |
|---|---|---|---|---|---|---|---|---|---|---|---|
| Spotted gum | *Corymbia maculata* | Myrtaceae | Tree | Regionally indigenous | | | | | VU | Single-stemmed tree 15-40 m tall. Bark smooth throughout, mottled grey over cream. Adult leaves concolorous, somewhat glossy and green. Buds and fruits occurring in groups of 3, grouped together to form larger semi-terminal clusters. Fruits broadly urn-shaped, 8-11 mm wide. Flowers Mar-Sep. | The natural distribution of Spotted Gum in Victoria is a very small population covering a few hectares near Mt Tara in the Mottle Range, south of Buchan. |
| Black-anther flax-lily | *Dianella revoluta* | Asphodelaceae | Lilioid | Locally indigenous | | | | | EN | Tufted perennial plants with stiffly erect linear leaves to 50 cm long. Dark green above, paler blue green below with recurved leaf margins, mid-rib smooth. Flowers blue to violet, anthers dark brown to black. Flowers held above foliage to 80 cm tall. Flowers Sep-Jan. | Occurs through a wide range of vegetation types from sea-level to subapls, and from mountain forests to mallee. |
| Ruby saltbush | *Enchylaena tomentosa* | Amaranthaceae | Shrub | Locally indigenous | | | | | EN | Prostate, spreading and sometimes erect shrub up to 1 m tall and wide. Small bluish green succulent leaves, branchlets downy. Flowers insignificant, hairy, greenish and axillary. Succulent green fruits ripen into yellow, orange and red berries. Flowers Sep-Jan. | Widespread in the north and north-west of Victoria in mallee scrub, in woodlands on heavy alluvial soils and on disturbed sites, occasional along the coast usually amongst rocks, and in rain-shadow areas to the north and west of Melbourne. |
| Weeping grass | *Microlaena stipoides* | Poaceae | Graminoid | Locally indigenous | | | | | NT | Sparsely tufted or shortly rhizomatous perennial grass that is highly variable in size, up to 1 m tall. Foliage smooth to rough flat leaves, top of sheath with few long hairs on ear-like lobes. Flowers narrow, weeping panicles to 18 cm long. Dark green to purplish spikelets. Flowers Oct-Mar. | Requires moist well-drained soils. Occurs in a wide variety of habitats, also gardens, lawns and pasture. |
| Common tussock-grass | *Poa labillardierei* | Poaceae | Graminoid | Locally indigenous | | | | | VU | Large, densely tufted perennial grass tussock with rough blue-green, flat bladed or tightly rolled (needle like) foliage, often with a sharp tip, to 80 cm tall. Branched flowering stems to 130 cm tall. Flowers Sep-Feb. | Common to a variety of habitats with good soil moisture, such as alluvial soils in riparian communities. |
| Cluster pomaderris | *Pomaderris racemosa* | Rhamnaceae | Shrub | Locally indigenous | | | | | EN | Slender shrub to 5 m tall. Leaves thin, dark green, ovate to broadly elliptic c. 2 cm long and 12 mm wide, upper surface glabrous, densely stellate-pubescent and pale green below. Flowers Oct-Dec. | Occurs on well-drained soils, scattered through sheltered forests and riparian scrub throughout South-west Victoria. |
| Kangaroo apple | *Solanum aviculare* | Solanaceae | Shrub | Locally indigenous | | | | | LC | Erect, woody shrub with angular stems to 4 m tall. Dark green leaves, variably lobed to 30 cm long and 15 cm wide. Some leaf shapes resemble a kangaroo paw. Upper leaves entire and lanceolate. Flowers violet with dark centres and deeply cut lobes. Succulent ovoid fruit, change from yellow-green to scarlet/ orange-red when ripe. Flowers Sep-Feb. | Found in a range of forest, woodland and swamp type habitats. Requires well drained soils, can grow in full sun to full shade. |

**Table S1 (Cont.)**. Detailed description of the 15 plant species that were part of the study.

| Common name | Species | Family | Growth form | Origin | Year 0 [2016] | Year 1 [2017] | Year 2 [2018] | Year 3 [2019] | IUCN cat. | General description | Distribution |
|---|---|---|---|---|---|---|---|---|---|---|---|
| Kangaroo grass | *Themeda triandra* | Poaceae | Graminoid | Locally indigenous | | | | | EN | Sprawling perennial tussock, stems sometimes reddish, older stems becoming branched. Foliage limp, flat or channelled, green to bluish leaves to 50 cm tall. Flower stems 10-25 cm long with red-brown triangular shaped spikelets, clustered along a droopy, leafy flowering stems. Flowers Dec-Feb. | Formerly, a dominant species over vast areas of the basalt plains of Victoria, occurs throughout Australia. Commonly associated with other grasses in woodland and grassland communities over a wide range of soil types and growing conditions, but not in permanently wet, very dry, saline or heavily shaded sites. |
| Tufted bluebell | *Wahlenbergia capillaris* | Campanulaceae | Forb | Locally indigenous | | | | | | Branching, erect perennial herb 15-50 cm tall with a thick tap root and few to many flowering stems from base. Leaves alternating up stem or clustered at base, 4-50 mm long, 0.5-6 mm wide. Flowers mauve to blue, sometimes white, tubular with 5 lobes, flowerheads on stalks above foliage. Flowers Oct-Mar. | This species is common throughout Australia in a range of grassland, woodland, forest and shrubland habitats. |

**Table S2**. Number of plant species included in each year of the study, including those that perished or were added each year.

| Plant species | Year 0 | Year 1 | | | | Year 2 | | | | Year 3 | | | |
|---|---|---|---|---|---|---|---|---|---|---|---|---|---|
| | Baseline | Perished | Added | Total | Turnover | Perished | Added | Total | Turnover | Perished | Added | Total | Turnover |
| All | 2 | 0 | 12 | 14 | 0.86 | 4 | 1 | 11 | 0.33 | 0 | 0 | 11 | 0.00 |
| Forbs | 0 | 0 | 1 | 1 | 1.00 | 0 | 0 | 1 | 0.00 | 0 | 0 | 1 | 0.00 |
| Lilioids | 0 | 0 | 1 | 1 | 1.00 | 0 | 0 | 1 | 0.00 | 0 | 0 | 1 | 0.00 |
| Graminoids | 1 | 0 | 3 | 4 | 0.75 | 1 | 0 | 3 | 0.25 | 0 | 0 | 3 | 0.00 |
| Shrubs | 0 | 0 | 5 | 5 | 1.00 | 3 | 1 | 3 | 0.67 | 0 | 0 | 3 | 0.00 |
| Trees | 1 | 0 | 2 | 3 | 0.67 | 0 | 0 | 3 | 0.00 | 0 | 0 | 3 | 0.00 |

**Table S3**. List of the 94 insect species that were recorded during the study. DET: Detritivore; HER: Herbivore; PRE: Predator; PAR: Parasitoid. All species are indigenous to the study area, excepting those marked with an *.

| Species/morphospecies | Common name | Family | DET | HER | PRE | PAR | Year 0 [2016] | Year 1 [2017] | Year 2 [2018] | Year 3 [2019] |
|---|---|---|---|---|---|---|---|---|---|---|
| **Hymenoptera \| Apocrita** | | | | | | | | | | |
| *Apocrita 5* | Apocrita 5 | | | | | ■ | | ■ | | |
| *Apocrita 33* | Apocrita 33 | | | | | ■ | | | | ■ |
| *Apocrita 36* | Apocrita 36 | | | | | ■ | | ■ | | |
| *Apocrita 37* | Apocrita 37 | | | | | ■ | | | ■ | |
| *Apocrita 40* | Apocrita 40 | | | | | ■ | | | ■ | |
| *Apocrita 44* | Apocrita 44 | | | | | ■ | | | ■ | |
| *Apocrita 46* | Apocrita 46 | | | | | ■ | | | ■ | |
| *Apocrita 48* | Apocrita 48 | | | | | ■ | | ■ | | ■ |
| *Apocrita 49* | Apocrita 49 | | | | | ■ | | ■ | | ■ |
| *Apocrita 51* | Apocrita 51 | | | | | ■ | | ■ | | ■ |
| *Apocrita 52* | Apocrita 52 | | | | | ■ | | ■ | | |
| *Apocrita 53* | Apocrita 53 | | | | | ■ | | ■ | | ■ |
| *Apocrita 54* | Apocrita 54 | | | | | ■ | | ■ | | ■ |
| *Apocrita 55* | Apocrita 55 | | | | | ■ | | | | ■ |
| *Apocrita 56* | Apocrita 56 | | | | | ■ | ■ | | | |
| *Apocrita 57* | Apocrita 57 | | | | | ■ | | ■ | ■ | ■ |
| *Apocrita 58* | Apocrita 58 | | | | | ■ | | | ■ | ■ |
| *Apocrita 59* | Apocrita 59 | | | | | ■ | | | ■ | ■ |
| *Apocrita 60* | Apocrita 60 | | | | | ■ | | | ■ | |
| *Apocrita 61* | Apocrita 61 | | | | | ■ | | ■ | ■ | ■ |
| *Apocrita 62* | Apocrita 62 | | | | | ■ | | | | ■ |
| *Apocrita 65* | Apocrita 65 | | | | | ■ | | | | ■ |
| *Apocrita 74* | Apocrita 74 | | | | | ■ | | | | ■ |
| *Apocrita 89* | Apocrita 89 | | | | | ■ | | | | ■ |
| *Apocrita 90* | Apocrita 90 | | | | | ■ | | | | ■ |
| *Apocrita 91* | Apocrita 91 | | | | | ■ | | | | ■ |

**Table S3 (Cont.)**. List of the 94 insect species that were recorded during the study. DET: Detritivore; HER: Herbivore; PRE: Predator; PAR: Parasitoid. All species are indigenous to the study area, excepting those marked with an *.

| | | | | | | | | | | |
|---|---|---|---|---|---|---|---|---|---|---|
| **Hymenoptera \| Apoidea \| Anthophila** | | | | | | | | | | |
| *Anthophila 1* | Anthophila 1 | | | ■ | | | | | | ■ |
| *Anthophila 3* | Anthophila 3 | | | ■ | | | | | | ■ |
| *Apis mellifera** | European honeybee | Apidae | | ■ | | | | | | ■ |
| **Diptera \| Brachycera** | | | | | | | | | | |
| *Brachycera 2* | Brachycera 2 | | ■ | | | | | ■ | | ■ |
| *Brachycera 7* | Brachycera 7 | | ■ | | | | | ■ | | ■ |
| *Brachycera 8* | Brachycera 8 | | ■ | | | | | ■ | | |
| *Brachycera 14* | Brachycera 14 | | ■ | | | | | ■ | ■ | ■ |
| *Brachycera 15* | Brachycera 15 | | ■ | | | | | ■ | | |
| *Brachycera 16* | Brachycera 16 | | ■ | | | | | ■ | | |
| *Brachycera 18* | Brachycera 18 | | ■ | | | | | ■ | ■ | |
| *Brachycera 19* | Brachycera 19 | | ■ | | | | | ■ | ■ | |
| *Brachycera 20* | Brachycera 20 | | ■ | | | | | ■ | ■ | ■ |
| *Brachycera 22* | Brachycera 22 | | ■ | | | | | | ■ | |
| *Brachycera 34* | Brachycera 34 | | ■ | | | | | ■ | | |
| *Brachycera 37* | Brachycera 37 | | ■ | | | | | ■ | | ■ |
| *Brachycera 38* | Brachycera 38 | | ■ | | | | | ■ | | ■ |
| *Brachycera 39* | Brachycera 39 | | ■ | | | | | ■ | | ■ |
| *Brachycera 40* | Brachycera 40 | | ■ | | | | | ■ | | ■ |
| *Brachycera 41* | Brachycera 41 | | ■ | | | | | ■ | | ■ |
| *Brachycera 42* | Brachycera 42 | | ■ | | | | | | ■ | ■ |
| *Brachycera 43* | Brachycera 43 | | ■ | | | | | ■ | | |
| *Brachycera 44* | Brachycera 44 | | ■ | | | | | ■ | | |
| *Tephritidae 3* | Tephritidae 3 | Tephritidae | | ■ | | | | ■ | | |
| **Coleoptera \| Chrysomeloidea** | | | | | | | | | | |
| *Chaetocnema sp.1* | Chaetocnema sp.1 | Chrysomelidae | | ■ | | | | ■ | | |
| *Diachus auratus* | Bronze leaf beetle | Chrysomelidae | | ■ | | | | | | ■ |
| *Hispellinus multispinosus* | Spiny leaf beetle | Chrysomelidae | | ■ | | | | | ■ | |

**Table S3 (Cont.)**. List of the 94 insect species that were recorded during the study. DET: Detritivore; HER: Herbivore; PRE: Predator; PAR: Parasitoid. All species are indigenous to the study area, excepting those marked with an *.

| | | | | | | | | | | |
|---|---|---|---|---|---|---|---|---|---|---|
| **Coleoptera \| Coccinelloidea** | | | | | | | | | | |
| *Coccinella transversalis* | Transverse ladybug | Coccinellidae | | | ■ | | | ■ | | |
| *Diomus pumilio* | Diomus pumilio | Coccinellidae | | | ■ | | | ■ | ■ | ■ |
| *Diomus sp. 1* | Diomus sp. 1 | Coccinellidae | | | ■ | | | | ■ | ■ |
| *Diomus sp. 2* | Diomus sp. 2 | Coccinellidae | | | ■ | | | ■ | | |
| *Diomus sp. 4* | Diomus sp. 4 | Coccinellidae | | | ■ | | | ■ | | |
| *Hippodamia variegata** | Spotted amber ladybug | Coccinellidae | | | ■ | | | | | ■ |
| *Serangium maculigerum* | Citrus whitefly ladybug | Coccinellidae | | | ■ | | | | ■ | ■ |
| **Hemiptera \| Heteroptera \| Coreoidea** | | | | | | | | | | |
| *Mutusca brevicornis* | Short-horned braod-headed bug | Alydidae | | ■ | | | | ■ | ■ | ■ |
| **Coleoptera \| Cucujoidea** | | | | | | | | | | |
| *Aethina concolor* | Sap beetle | Nitidulidae | ■ | | | | | ■ | ■ | |
| *Corticaria sp.1* | Minute brown scavenger beetle | Latriidae | ■ | | | | ■ | ■ | ■ | ■ |
| **Coleoptera \| Curculionoidea** | | | | | | | | | | |
| *Curculionidae 2* | Curculionidae 2 | Curculionidae | | ■ | | | | | ■ | |
| *Derelomini 1* | Derelomini 1 | Curculionidae | | ■ | | | | | | ■ |
| *Euciodes suturalis* | Fungus weevil | Anthribidae | ■ | | | | | ■ | ■ | |
| **Hemiptera \| Auchenorrhyncha \| Fulgoroidea** | | | | | | | | | | |
| *Anzora unicolor* | Grey planthopper | Flatidae | | ■ | | | | ■ | ■ | ■ |
| *Fulgoroidea 1* | Fulgoroidea 1 | | | ■ | | | | | | ■ |
| *Scolypopa australis* | Passionvine hopper | Ricaniidae | | ■ | | | | ■ | | |
| **Hemiptera \| Heteroptera \| Lygaeoidea** | | | | | | | | | | |
| *Nysius vinitor* | Rutherglen bug | Lygaeidae | | ■ | | | | ■ | ■ | ■ |
| *Stenophylla macreta* | Stenophylla macreta | Pachygronthidae | | ■ | | | | ■ | | ■ |
| **Coleoptera \| Membracoidea** | | | | | | | | | | |
| *Cicadellidae 4* | Cicadellidae 4 | Cicadellidae | | ■ | | | | ■ | ■ | ■ |
| *Cicadellidae 5* | Cicadellidae 5 | Cicadellidae | | ■ | | | | ■ | | |
| *Cicadellidae 6* | Cicadellidae 6 | Cicadellidae | | ■ | | | | | ■ | ■ |
| *Cicadellidae 7* | Cicadellidae 7 | Cicadellidae | | ■ | | | | ■ | | |
| *Erythroneurini 1* | Erythroneurini 1 | Cicadellidae | | ■ | | | | ■ | ■ | ■ |

**Table S3 (Cont.)**. List of the 94 insect species that were recorded during the study. DET: Detritivore; HER: Herbivore; PRE: Predator; PAR: Parasitoid. All species are indigenous to the study area, excepting those marked with an *.

| | | | | | | | | | | |
|---|---|---|---|---|---|---|---|---|---|---|
| **Hemiptera \| Heteroptera \| Miroidea** | | | | | | | | | | |
| *Coridromius sp. 1* | Coridromius sp. 1 | Miridae | | ■ | | | | ■ | | ■ |
| *Miridae 1* | Miridae 1 | Miridae | | ■ | | | | ■ | | ■ |
| *Miridae 2* | Miridae 2 | Miridae | | ■ | | | | ■ | | |
| *Miridae 3* | Miridae 3 | Miridae | | ■ | | | | ■ | | |
| *Miridae 5* | Miridae 5 | Miridae | | ■ | | | | | ■ | |
| *Miridae 7* | Miridae 7 | Miridae | | ■ | | | | ■ | ■ | |
| *Miridae 8* | Miridae 8 | Miridae | | ■ | | | | ■ | ■ | ■ |
| *Miridae 9* | Miridae 9 | Miridae | | ■ | | | | ■ | | ■ |
| *Sidnia kinbergi* | Australian crop mirid | Miridae | | ■ | | | | ■ | | ■ |
| *Thaumastocoridae 1* | Thaumastocoridae 1 | Thaumastocoridae | | ■ | | | | ■ | ■ | |
| **Naboidea** | | | | | | | | | | |
| *Nabis kinbergii* | Pacific damsel bug | Nabidae | | | ■ | | | ■ | | ■ |
| **Hemiptera \| Sternorryncha \| Psylloidea** | | | | | | | | | | |
| *Psyllidae 4* | Psyllidae 4 | Psyllidae | | ■ | | | | | ■ | ■ |
| *Psyllidae 5* | Psyllidae 5 | Psyllidae | | ■ | | | ■ | ■ | ■ | ■ |
| *Psyllidae 6* | Psyllidae 6 | Psyllidae | | ■ | | | ■ | ■ | ■ | ■ |
| *Psyllidae 7* | Psyllidae 7 | Psyllidae | | ■ | | | | ■ | | ■ |
| **Coleoptera \| Tenebrionoidea** | | | | | | | | | | |
| *Mordellidae 1* | Mordellidae 1 | Mordellidae | | ■ | | | | | ■ | |
| **Hymenoptera \| Vespoidea** | | | | | | | | | | |
| *Formicidae 1* | Formicidae 1 | Formicidae | | | ■ | | | | | ■ |
| *Formicidae 2* | Formicidae 2 | Formicidae | | | ■ | | ■ | ■ | ■ | ■ |
| *Vespula germanica** | European wasp | Vespidae | | | ■ | | | | | ■ |
| **Totals** | | | **22** | **35** | **11** | **26** | **5** | **60** | **41** | **59** |

**Table S4**. Number of insect species recorded in each year of the study, including those that went locally extinct (EXT) or colonised (COl) the site each year.

| Insect species | Year 0 | Year 1 | | | | Year 2 | | | | Year 3 | | | |
|---|---|---|---|---|---|---|---|---|---|---|---|---|---|
| | Baseline | EXT | COL | Total | Turnover | EXT | COL | Total | Turnover | EXT | COL | Total | Turnover |
| All | 5 | 1 | 56 | 60 | 0.93 | 40 | 21 | 41 | 0.75 | 17 | 35 | 59 | 0.68 |
| Detritivores | 2 | 0 | 19 | 21 | 0.90 | 11 | 3 | 13 | 0.58 | 5 | 7 | 15 | 0.60 |
| Herbivores | 3 | 0 | 21 | 24 | 0.88 | 13 | 6 | 17 | 0.63 | 7 | 15 | 25 | 0.69 |
| Predators | 1 | 0 | 5 | 6 | 0.83 | 4 | 2 | 4 | 0.75 | 0 | 4 | 8 | 0.50 |
| Parasitoids | 1 | 1 | 11 | 11 | 1.00 | 9 | 7 | 9 | 0.89 | 4 | 12 | 17 | 0.76 |

**Table S5**. Posterior estimates for species richness of indigenous insect species for the whole community and for each insect functional group as estimated under the hierarchical metacommunity model for baseline and greening action plant species for each year of the study.

| Insect species | Year 0 | | | | Year 1 | | | | Year 2 | | | | Year 3 | | | |
|---|---|---|---|---|---|---|---|---|---|---|---|---|---|---|---|---|
| | Mean | SD | 2.5% | 97.5% | Mean | SD | 2.5% | 97.5% | Mean | SD | 2.5% | 97.5% | Mean | SD | 2.5% | 97.5% |
| **All** | | | | | | | | | | | | | | | | |
| Baseline plant species | 3.227 | 0.261 | 3.000 | 3.500 | 10.150 | 1.057 | 8.000 | 12.000 | 5.339 | 0.675 | 4.000 | 6.500 | 5.562 | 0.650 | 4.000 | 6.500 |
| Greening action plant species | | | | | 15.810 | 0.865 | 14.167 | 17.500 | 14.376 | 1.004 | 12.444 | 16.333 | 23.469 | 1.302 | 20.889 | 26.222 |
| **Detritivores** | | | | | | | | | | | | | | | | |
| Baseline plant species | 1.500 | 0.000 | 1.500 | 1.500 | 3.135 | 0.581 | 2.000 | 4.000 | 2.729 | 0.469 | 2.000 | 3.500 | 3.662 | 0.587 | 2.500 | 4.500 |
| Greening action plant species | | | | | 6.327 | 0.672 | 5.167 | 7.667 | 4.747 | 0.440 | 3.889 | 5.556 | 8.645 | 0.517 | 7.444 | 9.556 |
| **Herbivores** | | | | | | | | | | | | | | | | |
| Baseline plant species | 2.000 | 0.000 | 2.000 | 2.000 | 5.089 | 0.716 | 4.000 | 6.500 | 3.070 | 0.423 | 2.000 | 3.500 | 3.408 | 0.523 | 2.500 | 4.000 |
| Greening action plant species | | | | | 8.208 | 0.594 | 7.182 | 9.455 | 7.273 | 0.602 | 6.000 | 8.222 | 10.378 | 0.884 | 8.556 | 11.778 |
| **Predators** | | | | | | | | | | | | | | | | |
| Baseline plant species | 1.000 | 0.000 | 1.000 | 1.000 | 3.001 | 0.562 | 2.000 | 3.500 | 1.990 | 0.380 | 1.500 | 2.500 | 1.777 | 0.302 | 1.000 | 2.000 |
| Greening action plant species | | | | | 2.137 | 0.261 | 1.600 | 2.500 | 1.638 | 0.316 | 1.143 | 2.143 | 4.434 | 0.414 | 3.444 | 4.889 |
| **Parasitoids** | | | | | | | | | | | | | | | | |
| Baseline plant species | 1.000 | 0.000 | 1.000 | 1.000 | 2.221 | 0.340 | 1.500 | 2.500 | 0.000 | 0.000 | 0.000 | 0.000 | 0.000 | 0.000 | 0.000 | 0.000 |
| Greening action plant species | | | | | 3.686 | 0.371 | 2.857 | 4.143 | 3.561 | 0.432 | 2.714 | 4.143 | 4.857 | 0.526 | 3.667 | 5.556 |

**Table S6**. Posterior estimates for the probability of occurrence, survival, and colonisation of indigenous insect species for the whole community and for each insect functional group as estimated under the multiseason site-occupancy model for each year of the study.

| Insect species | Year 0 | | | | Year 1 | | | | Year 2 | | | | Year 3 | | | |
|---|---|---|---|---|---|---|---|---|---|---|---|---|---|---|---|---|
| | Mean | SD | 2.5% | 97.5% | Mean | SD | 2.5% | 97.5% | Mean | SD | 2.5% | 97.5% | Mean | SD | 2.5% | 97.5% |
| **All** | | | | | | | | | | | | | | | | |
| Probability of occurrence | 0.257 | 0.124 | 0.057 | 0.532 | 0.859 | 0.091 | 0.631 | 0.981 | 0.881 | 0.080 | 0.690 | 0.985 | 0.874 | 0.081 | 0.687 | 0.983 |
| Probability of survival | | | | | 0.752 | 0.192 | 0.308 | 0.992 | 0.917 | 0.076 | 0.722 | 0.998 | 0.924 | 0.069 | 0.751 | 0.998 |
| Probability of colonisation | | | | | 0.894 | 0.094 | 0.654 | 0.997 | 0.665 | 0.240 | 0.151 | 0.987 | 0.497 | 0.293 | 0.022 | 0.977 |
| **Detritivores** | | | | | | | | | | | | | | | | |
| Probability of occurrence | 0.273 | 0.141 | 0.065 | 0.602 | 0.860 | 0.090 | 0.646 | 0.982 | 0.883 | 0.079 | 0.695 | 0.987 | 0.871 | 0.083 | 0.672 | 0.984 |
| Probability of survival | | | | | 0.756 | 0.191 | 0.306 | 0.993 | 0.919 | 0.074 | 0.719 | 0.998 | 0.920 | 0.073 | 0.728 | 0.997 |
| Probability of colonisation | | | | | 0.893 | 0.103 | 0.628 | 0.997 | 0.667 | 0.238 | 0.152 | 0.987 | 0.502 | 0.289 | 0.025 | 0.975 |
| **Herbivores** | | | | | | | | | | | | | | | | |
| Probability of occurrence | 0.272 | 0.137 | 0.064 | 0.581 | 0.861 | 0.088 | 0.649 | 0.982 | 0.880 | 0.081 | 0.676 | 0.986 | 0.873 | 0.084 | 0.667 | 0.983 |
| Probability of survival | | | | | 0.760 | 0.186 | 0.318 | 0.992 | 0.916 | 0.077 | 0.718 | 0.998 | 0.923 | 0.073 | 0.733 | 0.998 |
| Probability of colonisation | | | | | 0.895 | 0.095 | 0.645 | 0.997 | 0.662 | 0.237 | 0.162 | 0.987 | 0.499 | 0.287 | 0.027 | 0.975 |
| **Predators** | | | | | | | | | | | | | | | | |
| Probability of occurrence | 0.279 | 0.142 | 0.067 | 0.602 | 0.822 | 0.104 | 0.585 | 0.974 | 0.838 | 0.100 | 0.598 | 0.978 | 0.871 | 0.083 | 0.675 | 0.983 |
| Probability of survival | | | | | 0.757 | 0.189 | 0.320 | 0.992 | 0.884 | 0.099 | 0.628 | 0.996 | 0.920 | 0.076 | 0.716 | 0.998 |
| Probability of colonisation | | | | | 0.844 | 0.121 | 0.555 | 0.994 | 0.636 | 0.245 | 0.132 | 0.987 | 0.595 | 0.273 | 0.055 | 0.984 |
| **Parasitoids** | | | | | | | | | | | | | | | | |
| Probability of occurrence | 0.371 | 0.243 | 0.040 | 0.921 | 0.586 | 0.153 | 0.289 | 0.880 | 0.661 | 0.135 | 0.383 | 0.902 | 0.746 | 0.111 | 0.504 | 0.928 |
| Probability of survival | | | | | 0.426 | 0.259 | 0.022 | 0.932 | 0.729 | 0.167 | 0.361 | 0.979 | 0.879 | 0.105 | 0.612 | 0.996 |
| Probability of colonisation | | | | | 0.654 | 0.209 | 0.193 | 0.978 | 0.568 | 0.216 | 0.152 | 0.956 | 0.478 | 0.233 | 0.053 | 0.918 |

**Table S7**. Posterior estimates of network metrics for the community of indigenous insect species for each year of the study.

| Insect species | Year 0 | | | | Year 1 | | | | Year 2 | | | | Year 3 | | | |
|---|---|---|---|---|---|---|---|---|---|---|---|---|---|---|---|---|
| | Mean | SD | 2.5% | 97.5% | Mean | SD | 2.5% | 97.5% | Mean | SD | 2.5% | 97.5% | Mean | SD | 2.5% | 97.5% |
| Number of interactions | 3.515 | 0.942 | 1.948 | 5.644 | 60.766 | 3.917 | 53.448 | 68.503 | 34.034 | 3.330 | 27.822 | 40.799 | 51.399 | 4.076 | 43.671 | 59.728 |
| Interaction diversity | 1.043 | 0.183 | 0.680 | 1.407 | 3.603 | 0.182 | 3.242 | 3.967 | 3.267 | 0.208 | 2.854 | 3.690 | 3.584 | 0.212 | 3.157 | 4.000 |
| Interaction evenness | 0.825 | 0.060 | 0.706 | 0.946 | 0.742 | 0.063 | 0.616 | 0.863 | 0.708 | 0.072 | 0.560 | 0.850 | 0.762 | 0.072 | 0.618 | 0.904 |
| $H'2$ | | | | | 0.227 | 0.028 | 0.176 | 0.285 | 0.137 | 0.022 | 0.097 | 0.187 | 0.141 | 0.022 | 0.100 | 0.189 |
| $d'$ plants | | | | | 0.258 | 0.046 | 0.174 | 0.359 | 0.289 | 0.056 | 0.185 | 0.413 | 0.220 | 0.049 | 0.133 | 0.331 |
| $d'$ insects | | | | | 0.253 | 0.048 | 0.166 | 0.356 | 0.285 | 0.058 | 0.179 | 0.411 | 0.192 | 0.045 | 0.114 | 0.296 |
